# Alteration of Power Law Scaling of Spontaneous Brain Activity in Schizophrenia

**DOI:** 10.1101/2020.02.13.946657

**Authors:** Yi-Ju Lee, Su-Yun Huang, Ching-Po Lin, Shih-Jen Tsai, Albert C. Yang

## Abstract

Nonlinear dynamical analysis has been used to quantify the complexity of brain signal at temporal scales. Power law scaling is a well-validated method in physics that has been used to describe the complex nature of a system across different time scales. In this research, we investigated the change of power-law characteristics in a large-scale resting-state fMRI data of schizophrenia (N = 200) and healthy participants (N = 200) derived from Taiwan Aging and Mental Illness cohort. Fourier transform was used to determine the power spectral density (PSD) of resting-state fMRI signal. We estimated the power law scaling of PSD of resting-state fMRI signal by determining the slope of the regression line fitting to the log-log plot of PSD. The power law scaling represents the dynamical properties of resting-state fMRI signal ranging from noisy oscillation (e.g., white noise) to complex fluctuations (e.g., slope approaches −1). Linear regression model was used to assess the statistical difference in power law scaling between schizophrenia and healthy participants. The significant differences in power law scaling were found in six brain regions. Schizophrenia patients has significantly more positive power law scaling (i.e., frequency components become more homogenous) at four brain regions: left precuneus, left medial dorsal nucleus, right inferior frontal gyrus, and right middle temporal gyrus, compared with healthy participants. Additionally, schizophrenia exhibited less positive power law scaling (i.e., frequency components are more dominant at lower frequency range) in bilateral putamen. Significant correlations of power law scaling with the severity of psychosis were found in these identified brain areas in schizophrenia. These findings suggest that schizophrenia has abnormal brain signal complexity toward random patterns, which is linked to psychotic symptoms. The power law scaling analysis may serve as a novel functional brain imaging marker for evaluating patients with mental illness.

## Introduction

Interdisciplinary methods such as complexity science have been explored in recent years to understand the complex brain pathophysiology of schizophrenia. Complexity science has integrated the essences of physics, mathematics, and computational neuroscience, becoming the one of the emerging approaches to appraise the heterogeneous brain signal information. Despite typical reductionism epistemology, concepts such as entropy and fractal concepts of complexity science, as an evolutional paradigm for neuroscience, have been applied in human physiological system since 1990s.

In the central nervous system, complexity may reflect a system’s ability to adapt to the changing environment, which is often impaired in patients. Brain signals exhibiting in a random or regular manner are often observed in a pathological state, in contrast to the brain signals from a relatively healthy state (Goldberger et al., 2002). Yang and Tsai (2013) have proposed that the loss of brain complexity may result in mental dysfunction, and such decreased brain complexity was evidenced in the study of resting-state fMRI data (Xie et al., 2018; Yang et al., 2015). Defining functions of a physiological system with complexity parameters, from a prospect of frequency domain instead of the typical time domain, can provide diagnostic markers and prediction of therapeutic response in the clinic (Bystritsky et al., 2012; Fernandez et al., 2010; Manor and Lipsitz, 2013).

Additionally, the dynamical complexity of brain activity can be quantified by analyzing neural networks over a range of temporal scales. The aberrant neural structural or functional connectivity that indicates the deficit of information communication may play a role in mental illness, for example, in schizophrenia (Friston and Frith, 1995; Sokunbi et al., 2014). Such a perspective has inspired a number of studies in various neuropsychiatric disorders, such as sleep (Lo et al., 2002; Ma et al., 2017), mood disorders (Boettger et al., 2008; Voss et al., 2006), attention-deficit hyperactivity disorder (ADHD) (Ghassemi et al., 2012; Sokunbi et al., 2013), and autism spectrum disorder (Geschwind and Levitt, 2007; Wass, 2011).

The complexity of a given system can be described by three characteristics: nonlinearity, non-stationarity, and multiscale organization. For example, multiscale entropy analysis on brain functions was used to evaluate the temporal regularity or randomness of physiological signals (Liang et al., 2014; Yang et al., 2013; Yang et al., 2016). Yang et al. (2015) applied entropy analysis to resting-state fMRI data from patients with schizophrenia and documented that the decreased resting-state brain activity complexity can be characterized by increased regularity in the temporal lobe and increased randomness pattern in the frontal cortex, which explains the patients’ withdrawal behavior and auditory hallucinations. Nonlinear characteristics of brain signals were also reported in studies with various biomedical signal recording techniques, such as electroencephalogram (EEG) (Akar et al., 2016; Carlino et al., 2012; Cerquera et al., 2017; Di Lorenzo et al., 2015; Fernandez et al., 2013; Ibanez-Molina et al., 2018; Keshavan et al., 2004; Kim et al., 2000; Lee et al., 2008; Li et al., 2008; Raghavendra et al., 2009; Takahashi et al., 2010; Xiang et al., 2019; Yu et al., 2016) or magnetoencephalography (MEG) (Brookes et al., 2015; Fernandez et al., 2013; Fernandez et al., 2011; Kotini and Anninos, 2002; Robinson and Mandell, 2015).

Power-law distribution (or 1/f scaling in signal processing) is a ubiquitous principle in physics that describes the complex nature of a given system at multiple time scales and holds great potential to develop effective markers and to extract fundamental features from spatial-temporal neuroimaging data across levels. Power-law scaling has been observed in various neuronal system models. In 2000, an investigation on structural neural networks of all 302 neurons on Caenorhabditis elegans worm identified a power-law distribution of aging speeds in a whole nervous system (Amaral et al., 2000). In humans, the power-law phenomenon can be also observed across different levels of the human brain, such as neuronal firing rate (Buzsaki and Mizuseki, 2014), efficacy of synaptic transmission (Mizuseki and Buzsaki, 2013), channel density (Bullmore and Sporns, 2012), or neural circuit-level networks (Marković and Gros, 2014). In a system with power-law scaling, we could observe a skewed distribution of the relationship between two variables with a long right tail on a lognormal plot, which is often used to explain the complex nature of a given system at multiple time scales. The power law has been used to model various phenomena in nature and social sciences. In accordance with the loss of brain complexity hypothesis, brain activity in healthy state exhibits multiscale variability, which is a characteristic of power-law behavior, and schizophrenia brain could be associated with the breakdown of brain signal dynamics into regular or random patterns, for which its change in power law can be quantified rigorously via spectral analysis of resting-state fMRI signal. In this research, we hypothesized that spontaneous brain activity in schizophrenia may exhibit loss of power-law characteristics in the frequency domain compared to those observed in healthy volunteers. Therefore, we aimed to study the power-law scaling of resting-state fMRI signal in a large sample of schizophrenic patients and healthy subjects. The association between abnormal power law scaling and the Positive and Negative Syndrome Scale (PANSS) rating in schizophrenic patients is also examined.

## Material and Methods

### Participants

Functional brain imaging data of 200 age and sex matched schizophrenia patients (age mean = 43.56 ± 12.64; male = 49.5%) and 200 healthy subjects (age mean = 43.56±13.41; male = 49.5%), who were right-handed Han Chinese, were retrieved from Taiwan Aging and Mental Illness (TAMI) cohort. Diagnosis of schizophrenia was screened and confirmed by two psychiatrists based on criteria given in the Diagnostic and Statistical Manual of Mental Disorders (DSM–5). Written informed consent was obtained from all participants before the scanning sessions following the protocol for TAMI cohort approved by the review board at Taipei Veterans General Hospital, Taipei, Taiwan. All personal information and imaging data are de-identified for the subsequent analyses. It is worth mentioning that the TAMI cohort has recruited more than 1000 subjects so far, including a large sample of patients with healthy aging and patients covering major mental illness. All imaging data were acquired by the same 3.0T MRI Siemens Tim Trio machine with constant protocol at National Yang-Ming University.

### Data preprocessing

The resting-state fMRI image preprocessing was operated by DPARSF_V4.3_170105 (Data Processing Assistant for Resting-State fMRI) (Yan et al., 2016) and SPM12 (Statistical Parametric Mapping, Department of Imaging Neuroscience, London, UK) under MATLAB 2017a (Version 9.2). First, the first five of 200 data points for each participant were routinely discarded. Second, the functional images were motion-corrected and co-registered to the subject’s own anatomical images, followed by slice timing correction to the last slice. Third, the T1 image was realigned, and unified segmentation was used to normalize all functional images into standard Montreal Neurological Institute (MNI) space. Fourth, the BOLD signal was regressed out from both CSF and white matter space by Nuisance Covariates Regression. All participants in this study were selected from the TAMI cohort using the criteria of head motion that exhibited a maximum displacement of less than 1.5 mm at three axes and angular motion of less than 1.5° for each axis. We do not perform band-pass filtering for the purpose of power-law scaling estimation. Finally, Gaussian smoothing by SPM 12 with an 8 mm full width at half maximum (FWHM) was applied to all functional data to reduce noise from the spatial anatomical between-subject difference.

### Quantifying power-law scaling of resting-state fMRI signal

We first applied the Fourier Transform to the resting-state fMRI signal for each voxel to convert the time domain data into the frequency domain of the power spectrum (bin = 0.002 Hz), in order to quantify power-law scaling of the resting-state fMRI signal. Second, the data was visualized in a logarithm plot with the base equal to 10 on both axes to quantify the power spectrum across scales. Third, linear regression was deployed to derive a slope estimate, which was the scaling property of the given resting-state fMRI signal, to examine the distribution.

The dynamic signal of a system in a complex state exhibits a polynomial decrease, a typical power-law behavior in the frequency domain, which can be visualized as a negative slope close to minus one in the logarithm plot (left panel). The slope of the frequency scaling approaching minus one can provide evidence for the power-law behavior of the given resting-state fMRI signal. Therefore, such scaling analysis will be helpful for quantifying the complex dynamics of spontaneous brain activity, as shown in Fig. 1, in order to determine the state of brain activity at the complex state of 1/f power-law scaling behavior (left panel), reduced complexity as the slope of scaling becomes less steep (middle panel), and the brain signal to be an uncorrelated noise as the slope of scaling becomes flat (right panel).

**Figure 1.**
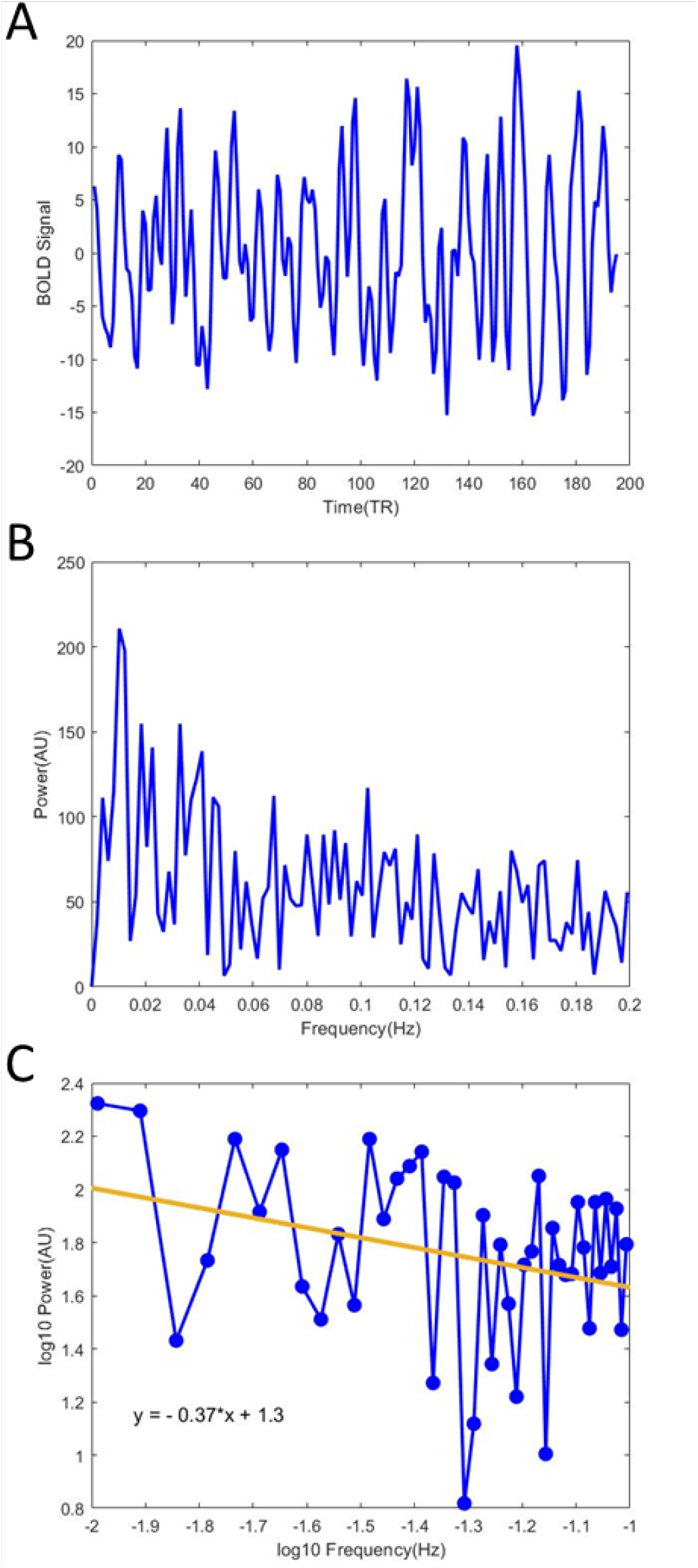
Quantifying power law scaling of brain BOLD signal in a voxel form real data. A. From a 30 years old healthy male’s resting-state fMRI image, we acquired a 4D BOLD time series data of a voxel at (24,14,36 of MNI coordinate) in left precuneus. B. A 3D power data was acquired after applying fast Fourier transform on the result of A. C. The signals were then transforming into a log-log plot. We use linear regression to acquire the slope of power law data, which indicates the power law scaling in a voxel. The power law scaling is calculated voxel-wise for both schizophrenic patients and healthy participants.

**Figure 2.**
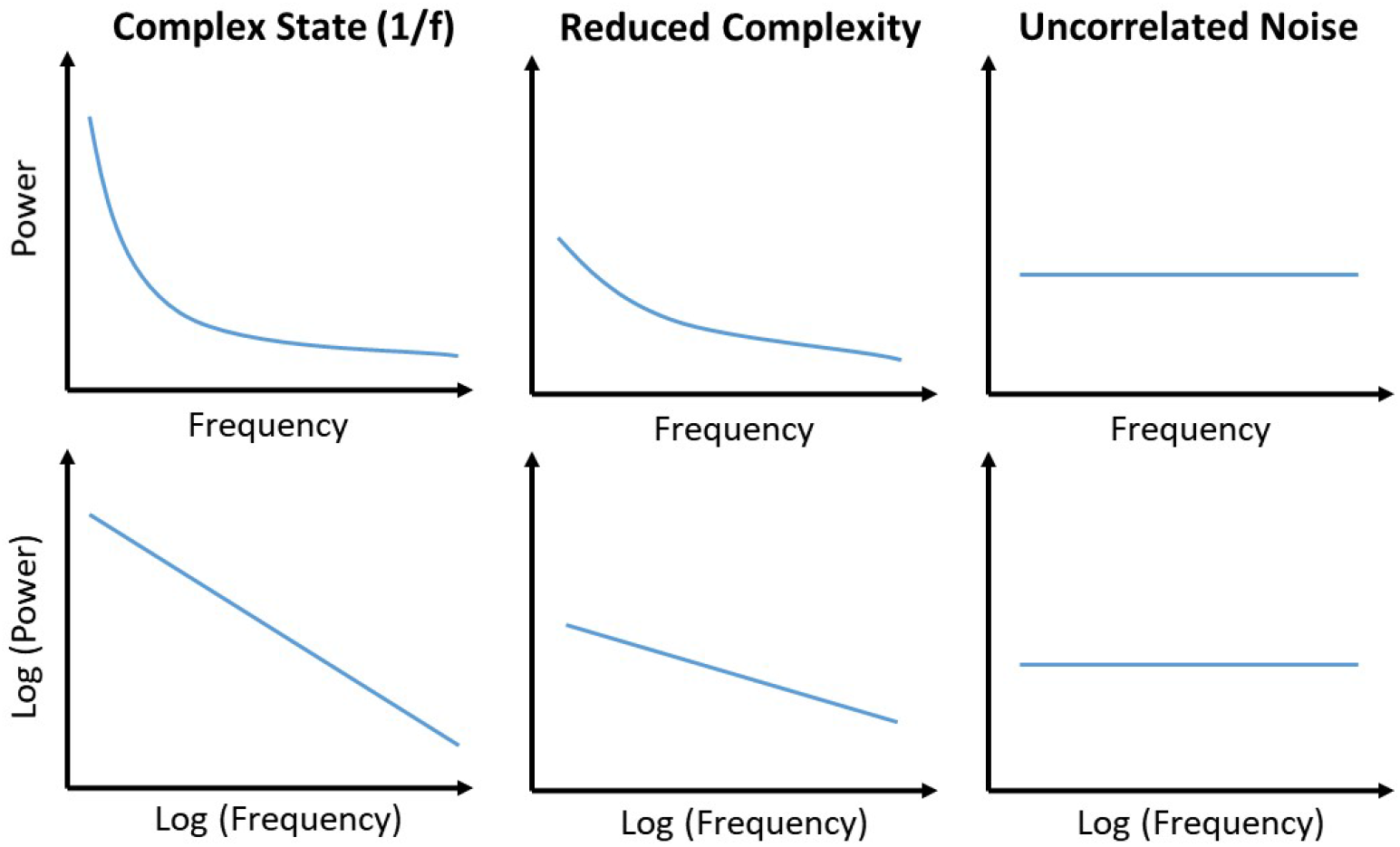
We visualize the power spectrum data onto a log-log plot with the base equal to 10 on both axes. Using linear regression, we can fit a regression line on the log-log plot and derive a slope estimate, which is the scaling property of the given resting-state fMRI signal. The slope of the frequency scaling approaching minus one would provide evidence for the power law behavior of the given resting-state fMRI signal. Therefore, such scaling analysis will be helpful for quantifying the complex dynamics of spontaneous brain activity in order to determine the state of brain activity at the complex state of 1/f power law scaling behavior (left panel), reduced complexity as the slope of scaling becomes less steep (middle panel), and the brain signal to be an uncorrelated noise as the slope of scaling becomes flat (right panel).

The power law distribution in the complex state has been observed in electrophysiological data (Gao, 2016; Tort et al., 2009; Voytek and Knight, 2015; Voytek et al., 2015), which can be linked to the physiological mechanism of neuronal communication. Such distribution has been also identified with electrophysiological data that is related to system dysfunction or increased difficulty in mental tasks (He, 2011, 2014; Miller et al., 2009; Podvalny et al., 2015; Pozzorini et al., 2013; Tinker and Velazquez, 2014; Usher et al., 1995; Voytek and Knight, 2015).

### Statistical analysis

Linear regression model is used to assess the between-group difference in voxel-wise power-law scaling data in gray matter to compare power-law scaling between schizophrenia patients and healthy adults. Both age and sex were used in the GLM as covariates. Significant brain clusters were reported if the nominal *P* value was less than 0.001 for any t-test (uncorrected) on a single voxel level with a cluster size greater than 10 voxels, and it was further corrected by family-wise error rate (FWE) methods at *P* values of less than 0.05 at the cluster level.

### Results

The linear regression model was used to compare the power-law scaling between schizophrenic patients and healthy adults (Table 1), with the extent threshold k = 35 voxels. Parametric images were assessed for cluster-wise significance using a cluster-defining threshold of FWE *p* < 0.05. The results reveal that schizophrenia patients, with the average onset of 28 years old and average duration of onset being 15 years, have significantly more positive power-law scaling and two more negative power-law scaling than healthy adults at four anatomical clusters.

**Table 1.** Coordinates of Clusters of Activation Showing Significantly Different Power Law Slope in Comparison Between Schizophrenic Patients and Healthy Adults. The table shows the results of t-test comparing the power law scaling between schizophrenia patients and healthy participants calculated by SPM12. The significant clusters reached of family-wise error rate (FWE) methods at P values of less than 0.05 at the cluster level with an extent threshold of 35 voxels. The power law behavior exhibits a negative slope in log-log plot. In compare to healthy participants, the brain activity of schizophrenia patients was found showing more positive slope (flatter, approach to 0) at four clusters: left precuneus, left thalamus, right middle frontal gyrus and right middle temporal gyrus, and more negative (steeper) slope in bilateral putamen.

| Region | *k* | *t* | Cluster level<br>*p*(corr) | Voxel level<br>*p*(FWE-cor) | *x* | *y* | *z* |
| --- | --- | --- | --- | --- | --- | --- | --- |
| Schizophrenia Patients > Healthy Adults |  |  |  |  |  |  |  |
| Left Precuneus | 17555 | 7.72 | 0.000 | 0.000 | -18 | -66 | 18 |
| Left Middle Occipital Gyrus |  | 6.97 | - | 0.000 | -21 | -93 | 18 |
| Left Middle Occipital Gyrus |  | 6.87 | - | 0.000 | -33 | -96 | 3 |
| Left Thalamus | 183 | 5.99 | 0.000 | 0.000 | -6 | -15 | 3 |
| Right Thalamus |  | 5.80 | - | 0.000 | 12 | -15 | 9 |
| Right Middle Frontal Gyrus | 160 | 4.26 | 0.000 | - | 42 | 21 | 30 |
| Right Middle Temporal Gyrus | 48 | 3.93 | 0.003 | - | 51 | -39 | -3 |
| Healthy Adults > Schizophrenia Patients Left |  |  |  |  |  |  |  |
| Right Putamen | 60 | -3.11 | 0.001 | - | 18 | 6 | -3 |
| Left Putamen | 44 | -3.11 | 0.005 | - | -18 | 9 | 3 |

Note: all clusters reached a significance level of $p < .001$ and an extent threshold of 35 voxels. For each cluster, coordinates are given for the maximally activated voxel and up to two local maxima *k*, number of voxels in cluster; *x*, *y*, *z*, coordinates (mm)

The four more positive clusters included left precuneus (*k* = 17,555; peak coordinate (mm) = −18, −66, 18; T = 7.72), with sub-cluster at left middle occipital gyrus (peak coordinate (mm) = −21,−93,18; T = 6.97), left medial dorsal nucleus (*k* = 183; peak coordinate (mm) = −6,−15, 3; T=5.99), right inferior frontal gyrus (*k* = 160; peak coordinate (mm) = 42, 21, 30; T = 4.26), and right middle temporal gyrus (*k* = 48; peak coordinate (mm) = 51, −39, −3; T = 3.93). All these four clusters have *p(FWE-cor)* < 0.001 at voxel level. The equivalent *k =* 37 of left Insula reached threshold of *k* = 35, however, at cluster level, the uncorrected *p* remained marginally significant. On the other hand, healthy adults demonstrated significantly higher power-law scaling than schizophrenia patients in two regions: right putamen (*k* = 60; peak coordinate (mm) = 18, 6, −3; T = −3.11) and left putamen (*k* = 44; peak coordinate (mm) = −18, 9, 3; T = −3.11).

Two anatomical clusters presenting more negative power-law scaling in schizophrenia patients are right putamen (*k* = 60; peak coordinate (mm) = 18, 6, −3; T = −3.11) and left putamen (*k* = 44; peak coordinate (mm) = −18, 9, 3; T = −3.11). These two clusters have *p (FRD-cor)* < 0.001 at the voxel level. The power law slope of these regions based on AAL90 template were calculated as Table 2.

**Table 2.** The Power Law Scaling of Identified Brain Regions We derived the power law scaling in identified AAL90 brain regions, where significant differences were found between schizophrenic patients and healthy subjects by t-test. The value indicates the mean and standard deviation of power law slope in log-log plot. In left precuneus, left thalamus, right middle frontal gyrus and right middle temporal gyrus, power law scaling in healthy participants exhibit more negative slope (closer to −1). In addition, the healthy participants show flatter slope (close to 0) in bilateral putamen in compare to schizophrenia patients.

The Power Law Scaling of Identified Brain Regions
| Brain Regions | Healthy |  | Schizophrenia |  |
| --- | --- | --- | --- | --- |
|  | mean | *S. D.* | mean | *S. D.* |
| Left Precuneus | -1.05 | 0.20 | -0.95 | 0.22 |
| Left Thalamus | -0.44 | 0.18 | -0.38 | 0.17 |
| Right Middle Frontal Gyrus | -0.78 | 0.16 | -0.72 | 0.18 |
| Right Middle Temporal Gyrus | -0.84 | 0.18 | -0.75 | 0.19 |
| Right Putamen | -0.35 | 0.11 | -0.38 | 0.13 |
| Left Putamen | -0.37 | 0.11 | -0.41 | 0.14 |

The linear regression model was used to compare the power-law scaling change between schizophrenic patient and healthy participants. The result visualization is shown in Figure 3, presenting the distribution of voxel-wise t-value across the brain. In Figure 3, red color indicates positive t value whereas blue color represents more negative t value. The swift of power-law scaling of resting-state fMRI signal indicates the state change in the brain system.

**Figure 3.**
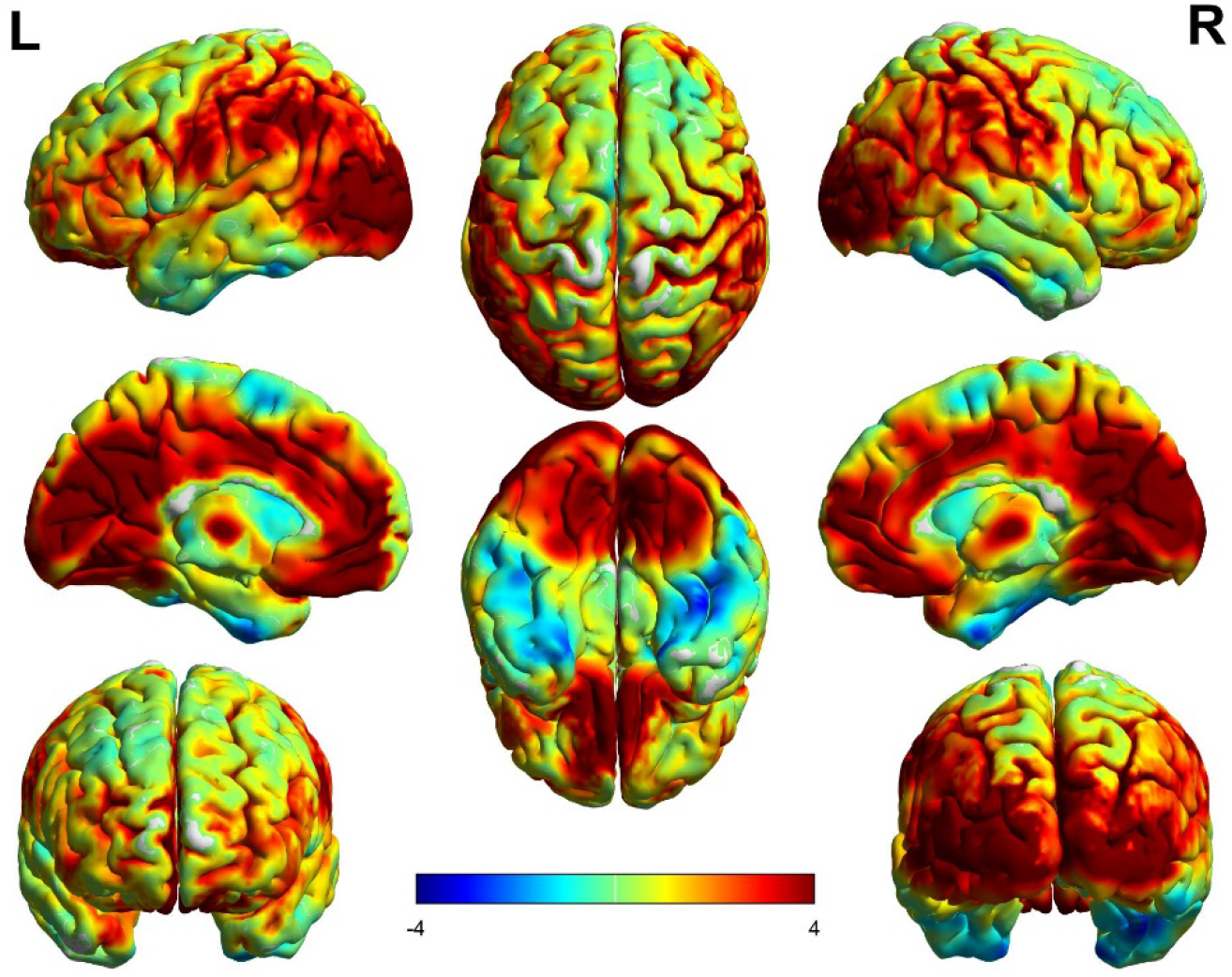
Visualization of difference in power law scaling of resting-state fMRI signal observed between schizophrenia patient and healthy participants. The power law scaling of resting-state fMRI signal was calculated for all voxels. Linear regression model was used to assess the between-group difference in power law scaling. Significant differences were found in multiple brain regions, including left precuneus, left thalamus, right middle frontal gyrus, right middle temporal gyrus, and bilateral putamen. The color bar indicates the strength of t-value that positive t-value indicates flatter slope of power law scaling of resting-state fMRI signal in schizophrenia, whereas negative t-value represents steeper slope of power law scaling in schizophrenia.

The Pearson correlation between the abnormal power-law scaling and score of PANSS is calculated. Interestingly, significant correlations with p-value < 0.05 were found in the key regions where the slope of power-law scaling was more positive in schizophrenic patients. The positive correlation was found between the power-law scaling slope in left precuneus and score of item G5 (mannerisms & posturing, *r* = 0.15, *p* = 0.036) and left thalamus and score of item N3 (passive/apathetic social withdrawal, *r* = 0.17, *p* = 0.017). Negative correlations were found between right middle frontal gyrus and score of item P4 (*r* = −0.15, *p* = 0.031) and right middle temporal gyrus and score of item P4 (excitement, *r* = −0.153, *p* = 0.032).

Pearson’s correlation is used to evaluate the relationship between the dosage of antipsychotic drugs and power-law scaling of resting-state fMRI signals in the identified brain regions. The antipsychotic dosage was transformed into Chlorpromazine (CPZ) equivalence dosage based on empirical studies (Andreasen et al., 2010; Danivas and Venkatasubramanian, 2013; Gardner et al., 2010). There was no significant correlation between CPZ dosage and the power-law scaling in six brain regions presented in Table 1.

## Discussion

The key findings in this study includes the following: (1) The difference in power-law scaling behavior in different anatomical regions indicates that schizophrenia is associated with the abnormal complexity of spontaneous brain activity in gray matter; and (2) the identified brain regions with abnormal complexity found in schizophrenia patients are correlated to psychotic symptoms, such as abnormal mannerism and posturing, excitement, and passive or apathetic social withdrawal. Collectively, these findings may support the Loss of Brain Complexity Hypothesis (Yang and Tsai, 2013) and suggest the reduced complex brain activity as one of the neurobiological mechanisms in schizophrenia. Additionally, the logarithmic transformation of the power spectrum provides a useful representation to observe brain signal dynamics, allowing us to examine the complexity change in the frequency domain. The pathological change in functional brain complexity suggests that power-law scaling is a ubiquitous principle of governing brain signal dynamics, which can potentially serve as a brain-based biomarker for schizophrenia.

The identified gray matter regions with abnormal power-law scaling are implicated in schizophrenia. The abnormal power-law scaling in the precuneus and left occipital gyrus was found to correlate with clinical symptoms in the present study. These brain regions are related to the visual function that is responsible for the interpretation of the images. The lesion in this area is evidenced in the study of visual hallucinations (McCarthy-Jones et al., 2017; Oertel et al., 2007; Stephan-Otto et al., 2017). Furthermore, decreased amplitudes of low-frequency oscillations (Hoptman et al., 2010) of resting-state fMRI and reduced negative connectivity (Zhou et al., 2007) in these areas were observed in schizophrenia.

The left medial dorsal nucleus in the thalamus is critical in attention and abstract thinking and is implicated in the sensory information processing (Welsh et al., 2010). The functional abnormalities in this region in schizophrenia may be induced by synaptic degeneration (Blennow et al., 1996), neuro-transmitter dysregulation such as glutamate and glutamine (Théberge et al., 2002), or metabolism difference revealed by positron emission tomography (Hazlett et al., 1999).

The right inferior frontal gyrus is the crucial area in the executive control and response inhibition. The anterior end is found to be related to cognition-related functions such as reasoning and social-cognitive processes, and the posterior end of this gyrus is related to action-related functions (Hampshire et al., 2010; Hartwigsen et al., 2018). For schizophrenic patients, the deficit of anatomical and functional brain connectivity associated with this region is found correlated with impairment of stop-signal inhibition (Aron et al., 2003; Swick et al., 2008), attentional control (Hampshire et al., 2010; Matzke et al., 2017), and auditory and verbal hallucinations (Lawrie et al., 2002; Lennox et al., 2000; Sommer et al., 2008),.

The mechanism that lead to the abnormal power law scaling of brain activity requires discussion.

Various psychiatric disorders, such as schizophrenia, depression, and autism spectrum disorder, are found to associate with the deficit in right middle temporal gyrus. Genetic factors which lead to the abnormal gray matter volume of this region have been suggested. (Cauda et al., 2011; Glahn et al., 2010; Krug et al., 2011). The reduced gray matter volume in this area is involved in auditory hallucinations (Kuroki et al., 2006; Onitsuka, 2004). Specifically, the G72 rs1421292 polymorphism may be associated with verbal function in schizophrenia (Donohoe et al., 2007; Krug et al., 2011).

On the other hand, power-law scaling in bilateral putamen was found to have a steeper slope (more negative) in schizophrenia compared to healthy volunteers (Figure 4). This may implicate in overactivity of dopaminergic neurons in this area in schizophrenia (Grace, 2016; McCutcheon et al., 2019). In Luo et al. (2019)’s study, SLC39A8 is a genetic variant involved in the anatomical development of putamen. Furthermore, this genetic variant altering putamen developmental trajectories contributes to the neuronal ion transport deficit in schizophrenic patients. The function of dorsal-ventral distribution has been studied, where putamen plays the crucial rule in anti-psychotic acting by inhibiting the uptake of dopamine from nigrostriatal tracts (McCutcheon et al., 2019). By investigating the aberrant mechanism in bilateral putamen of schizophrenic brain, it is possible that the more negative power scaling in putamen may indicate an overactivation in these dopaminergic regions, contributing to the cognitive symptom severity of schizophrenia.

**Figure 4.**
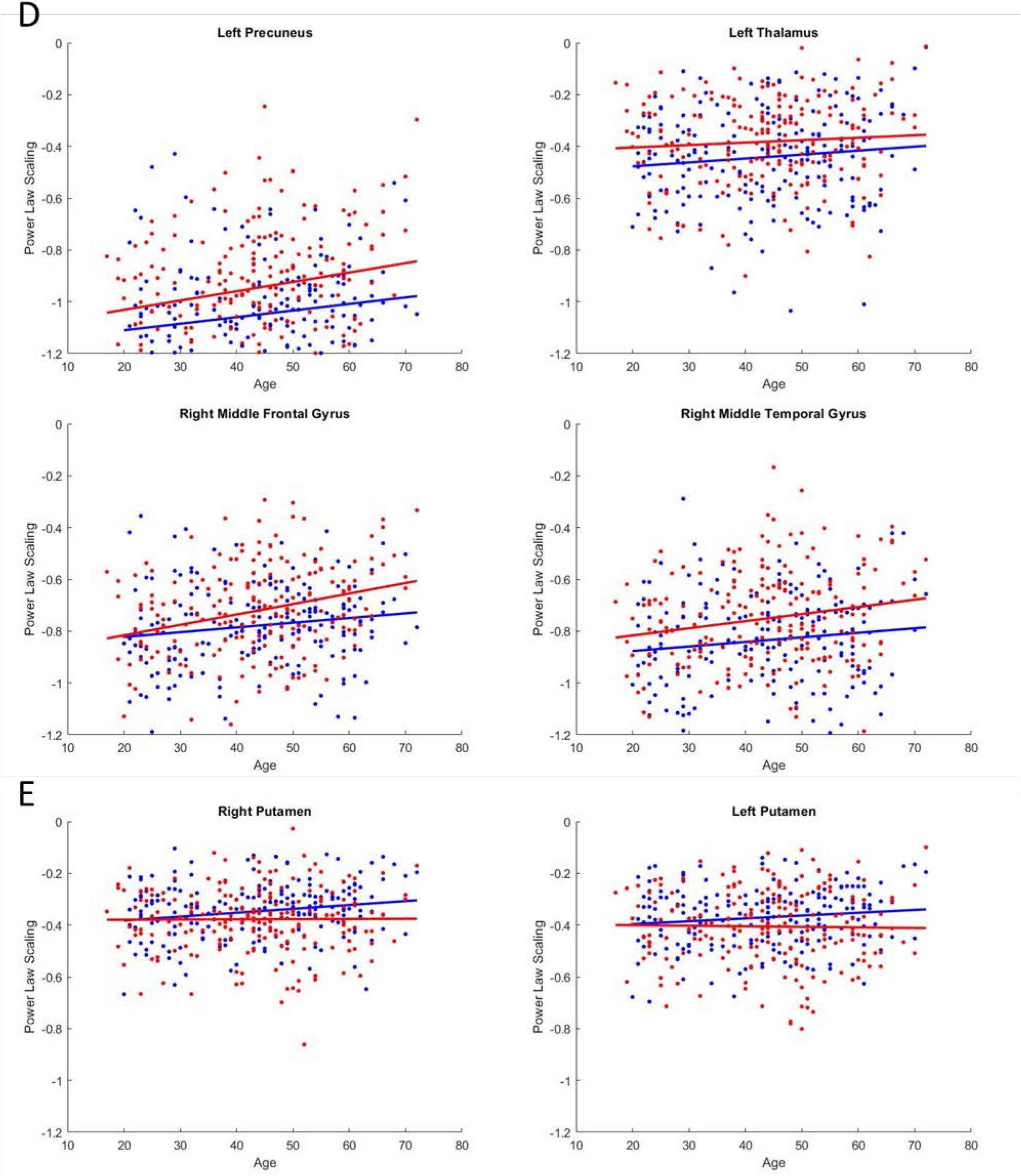
The power law scaling of identified brain regions for each participant were presented. Red indicates the power law scaling of schizophrenic patients. Blue indicates the power law scaling data of healthy participants. The regression line was applied for both datasets. In D, left precuneus, left thalamus, right middle frontal gyrus and right middle temporal gyrus, the power law scaling in healthy participants were closer to −1 across age. On the other hand, the figure E shows the power law scaling of schizophrenic patients approaching to −1 in bilateral putamen across age.

Our study findings indicate that spectral density of resting-state fMRI signals in healthy volunteers exhibits a higher power in lower frequency bins and lower power in higher frequency bins compared to schizophrenic patients, and such scaling behavior is more close to 1/f characteristics in healthy volunteers than that observed in schizophrenia. These findings suggest that schizophrenia may have a loss of 1/f brain signal dynamics, which indicates a possible loss of scale-free brain signal dynamics.

From the perspective of complexity science, signals observed in a dysfunctional system may demonstrate random or regular behavior. Change in power-law scaling toward a flat slope indicates a loss of multi-scale complexity, which is possibly associated with a lack of thinking or behavioral flexibility commonly observed in psychotic patients. Additionally, flattened power-law scaling observed in schizophrenia may also indicate an increased noise of information flow in the neuronal systems, which may be associated with abnormal structural or functional connectivity.

Estimating the interdependencies between different brain regions is crucial to understand the patterns of brain networks. Adapting from mathematical theory, neurobiologists use topology to represent the network behavior of neuronal orchestration. Interestingly, the scale-free behavior observed in one-dimensional time series of resting-state fMRI signals may be related to small-world network identified in the functional brain connectomes. In contrast to the random network topology, which the number of links of each node follows a normal distribution, the network topology in a complex system usually demonstrates a connection pattern following power-law distribution, enabling the complex network to be balanced between cost and efficiency (Bullmore and Sporns, 2012). Evidence from diffusion tensor image studies has suggested the deficit of structural connectivity in schizophrenic patients, indicating that the deficit of information flow in the brain network might be the pathophysiology underlying schizophrenic symptoms (Lener et al., 2014). A further study is needed to explore the relationship between power-law scaling in one-dimensional fMRI time series and small-world property derived from functional brain connectomes. Furthermore, causality methods can be also deployed to quantify the directionality and information flow between brain regions.

Complexity theory has been introduced to medical fields for a long time, such as cardiovascular physiology or human behavior. We have demonstrated that resting-state fMRI signals in schizophrenia exhibited two ways of abnormal complexity toward either regularity or randomness using an entropy approach (Yang et al., 2015). In this study, we further documented the use of frequency domain analysis to quantify the power-law scaling behavior of resting-state fMRI signals, showing loss of power scaling (i.e., toward randomness) in fMRI signals of schizophrenic patients, which is consistent with our previous report using the entropy approach (Hager et al., 2017; Yang et al., 2015). The analysis of power-law scaling in fMRI data is a new trend, especially in brain image studies. Overall, this emerging paradigm of complexity-based quantification has the potential to advance the understanding of brain functions and provide an objective imaging marker for evaluating patients with mental illness.

This study has several limitations. First, the frequency resolution of the power spectrum may be limited by the use of Fourier transform due to relatively short resting-fMRI time series. A better frequency resolution may be achieved by increasing the scanning time or the use of sophisticated time-frequency analysis, such as Hilbert-Huang transform. Second, whether changes in power-law slope in gray matter regions are related to alterations of neuronal dynamics requires further validation. Typically, resting-state fMRI signal between 0.01 and 0.1 Hz in gray matter regions is considered to be related to the neuronal activity via the mechanisms of neurovascular coupling. We have previously found that the cognitive-related function may be associated with a narrower band between 0.045–0.087 Hz (Yang et al., 2018). The power-law dynamics presented in this study could be refined in future study. Third, linear regression of the power spectrum may overlook certain dynamics that could be studied further with other methods of scaling dynamics, such as detrended fluctuation analysis (DFA). However, DFA requires long-time series data and may not be applicable to this study. A future study with a longer scanning time or a higher sampling rate may provide more information about brain signal dynamics.

## Conclusions

The application of complexity science in neuroscience has overcome common bias and concerns of typical statistic methods. It has presented insights that transcend traditional biological knowledge. The power-law phenomenon has emerged based on the integration of various biological mechanisms across temporal and spatial levels, supporting the discovery of nonlinear behavior of neuronal activity in the human central nervous system. Based on the frequency domain, in this research, power scaling plays a role to differentiate schizophrenic and healthy brain by analyzing resting-state brain imaging signals. Moreover, the key regions that show a significant difference in power scaling, such as left precuneus, left medial dorsal nucleus, inferior frontal gyrus, and bilateral putamen, correspond with clinic observations. In the near future, power-law analysis has great potential to offer markers in clinics and help to differentiate psychiatric disorders with different populations as well as suggesting diagnostic decisions and monitoring therapeutic processes.

## Acknowledgements

This work was supported by the Brain Research Center, National Yang-Ming University from The Featured Areas Research Center Program within the framework of the Higher Education Sprout Project by the Ministry of Education (MOE) and the Ministry of Science and Technology (MOST) of Taiwan (grant MOST 108-2634-F-075-002). Dr. Yang was supported by Mt. Jade Young Scholar Award from MOE.

